# Electrochemical measurement of acetylcholine in the dorsolateral prefrontal cortex: A technical report

**DOI:** 10.1101/120691

**Authors:** Devavrat Vartak, Chris van der Togt, Bram van Vugt, Pieter R. Roelfsema

**Affiliations:** Department of Vision & Cognition, Netherlands Institute for Neuroscience, Meibergdreef 47, 1105 BA, Amsterdam, The Netherlands.; Department of Integrative Neurophysiology, Center for Neurogenomics and Cognitive Research, VU University, Amsterdam, The Netherlands.; Psychiatry Department, Academic Medical Center, Amsterdam, The Netherlands

## Abstract

Ever since the discovery of acetylcholine in 1913, its role as neuromodulator has been extensively studied in a variety of model systems. These previous studies revealed that acetylcholine is of critical importance for several cognitive functions including attention, learning and memory. In spite of these previous findings, it has proven difficult to determine the amount of acetylcholine that is released during cognitive tasks with sub-second temporal resolution. One method that might be used to measure acetylcholine release is the use of an enzyme-coupled amperometric sensor, which has been suggested to measure acetylcholine with high sensitivity, selectivity and relatively high temporal resolution (< 1 second). In the present study, we have tried to adapt the technique developed in the rodent model system by Parikh and colleagues^1,2^ for use in non-human primates. We aimed to measure in-vivo levels of acetylcholine in the macaque dorsolateral prefrontal cortex while the monkey performed an attention demanding curve-tracing task^3,4^. We report that our attempts to measure acetylcholine using amperometry in an awake behaving macaque monkey proved difficult and tedious and that our results are inconsistent and prone to noise. In the discussion, we will outline the challenges that will need to be addressed to use this technique in non-human primates and hope that our observations inspire solutions to help future research on the role of this important neurotransmitter.

## Introduction

Acetylcholine was first discovered by Henry Dale^5^ in 1913. Ever since the discovery of acetylcholine, its role in the central nervous system has been studied extensively in a variety of model systems. The cholinergic projections from the nucleus basalis of Meynert innervate various areas of the cerebral cortex in mammals, suggesting that the cholinergic system and its innervations in the cortex directly and indirectly affect a wide range of cognitive functions^6,7^.

A large number of neurophysiological and pharmacological studies implicate the cortical cholinergic system in attentional performance. Indeed, attention demanding tasks recruit the basal forebrain cholinergic projections^8–11^. However, other cognitive functions, including reward processing^12^, learning^13^ and memory processing^14–16^ have also been linked to the release of acetylcholine. Yet other studies have suggested an important role of this neuromodulator in the development of the visual cortex and also in cortical plasticity in adult animals^17–20^.

Important insights into the function of acetylcholine can be derived from measurements of the amount of acetylcholine released during various cognitive tasks. Over the years, multiple techniques have been used to measure levels of acetylcholine in the brain^21,22^. A popular technique to measure the levels of acetylcholine has been microdialysis^23–27^. This technique involves the insertion of a probe which is used to sample the extracellular fluid containing the molecule(s) of interest via diffusion across a semi-permeable membrane. The collected fluid (microdialysate) is then analysed, for example, using chromatographic techniques coupled with electrochemical detection^28,29^ or spectrometric techniques^30–32^ to determine the concentration of the molecule of interest. Although microdialysis has been an insightful method to understand the regulation^27^ and quantification^26^ of acetylcholine in the brain, the temporal resolution of this technique is low, requiring several seconds to minutes per sample. It cannot be used to quantify rapid fluctuations in acetylcholine levels occurring during individual trials of a cognitive task. Shifts of attention, for example, occur on a time-scale of tens of milliseconds and changes in the concentration of acetylcholine cannot be detected by microdialysis at this time scale.

Several efforts have been made to develop techniques which allow the measurement of neuro-modulators at a faster temporal resolution. One such method is the use of amperometry/voltammetry using an enzyme catalyst. The electrode’s recording surface, usually an inert metal such as platinum (Pt), can oxidize or reduce compounds of interest when a potential is applied relative to a reference electrode (e.g. a silver-silver chloride (Ag/AgCl) electrode). If the potential at the electrode surface is sufficient, molecules in the vicinity of the surface of the electrode are either oxidized or reduced, depending upon their intrinsic electrochemical properties, at a rate that is proportional to their concentration. As the process of oxidation/reduction is quick, it causes changes in the current with a sub-second time resolution. The crucial step in these electrochemical methods is the oxidation/reduction reaction, which takes place at the recording surface. However, many biological substances do not undergo spontaneous oxidation/reduction on the platinum recording surface and an extra step is required to facilitate this process. Some methods use a catalyst such as a biological enzyme that converts the molecule of interest into an electro-active substance that will react on the recording surface. This method can be combined with the application of ‘exclusion layers’ using special polymers that prevent the oxidation/reduction of other substances, hence improving the selectivity of the measurement.

Parikh and colleagues^1^ used an amperometric method to detect rapid changes in the levels of acetylcholine in the cortex of rats. They used a bio-sensor developed by Burmeister and colleagues^33^. The key components of the bio-sensor are the ceramic sensor (Fig. 1b,c), which has platinum electrode contacts. Some substances undergo oxidation/reduction on the Pt recording surface when an appropriate voltage is applied with a potentiostat, but acetylcholine does not. However, when acetylcholine is released in the synaptic cleft, it is cleaved into choline by the enzyme acetylcholinesterase, which is present in the synaptic cleft in high concentrations. The choline oxidase enzyme then converts the newly formed choline into hydrogen peroxide and betaine. Hydrogen peroxide undergoes oxidation (at ∼0.7 V, pH of 7.0), thus generating a current across the electrode when an appropriate voltage is applied with a potentiostat. In order to increase the specificity for choline, an ‘exclusion layer’ such as nafion or m–(1,3)–phenylenediamine (mPD) is coated on the electrode, preventing larger molecules (or molecules with a particular charge) from reaching the Pt contact sites. Figure 1a illustrates the multisite microelectrode array, the coating of the sites and the steps involved in the conversion of choline to hydrogen peroxide. Another study by Parikh and colleagues^2^ demonstrated the use of this amperometric technique to measure acetylcholine levels for cue detection in the frontal cortex of rats. The study reported that acetylcholine release is stronger when the rat detects the cue than if the animal misses it. Furthermore, cortical acetylcholine appeared to be regulated at various time scales, from seconds to several minutes.

**Figure 1.**
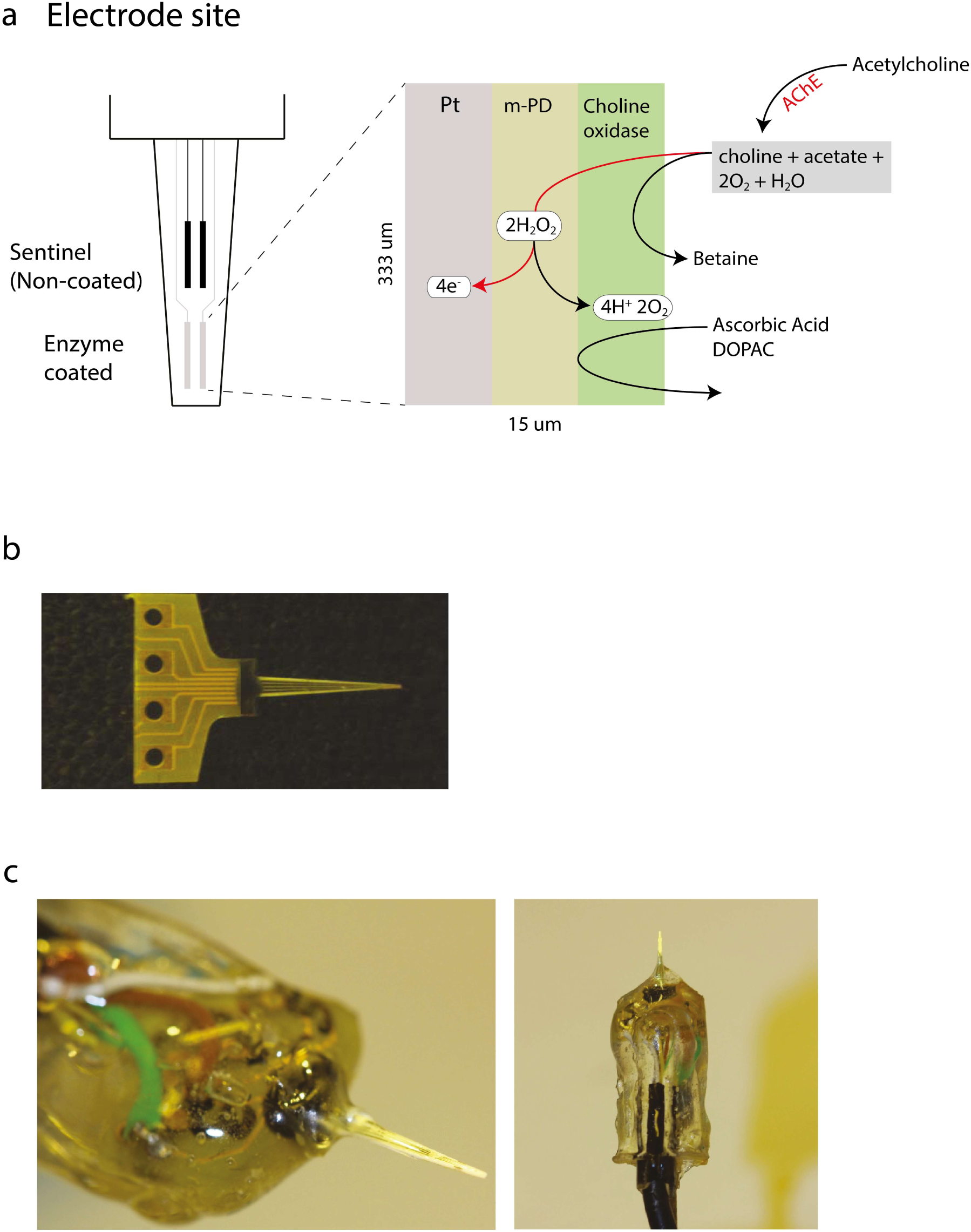
Electrode design. (a) Schematic of the electrode design. The recording surface and the reactions taking place in the different layers of the electrode. The Pt electrode was coated with an exclusion layer (m-PD) and an enzyme layer (choline oxidase). (b) Picture of the S2 electrode. (c) Picture the electrode that had been prepared for recording. Note the thick layer of insulation material that helped in reducing the noise level.

We here aimed to measure acetylcholine release during an attention task in an awake behaving macaque monkey. A robust paradigm that requires shifts of attention is the curve tracing task^3,4^ (Fig. 2a). The monkey fixates on a dot and sees two curves, one of which is connected to the fixation dot. One of these curves is the target curve connecting the fixation dot to a larger circle, which is the target for an eye movement (‘t’ in Fig. 2a). The monkey has to maintain fixation while deciding which one of the two curves is the target curve and the distractor curve. The monkey then indicates his decision by making a saccade to the endpoint of the target curve.

**Figure 2.**
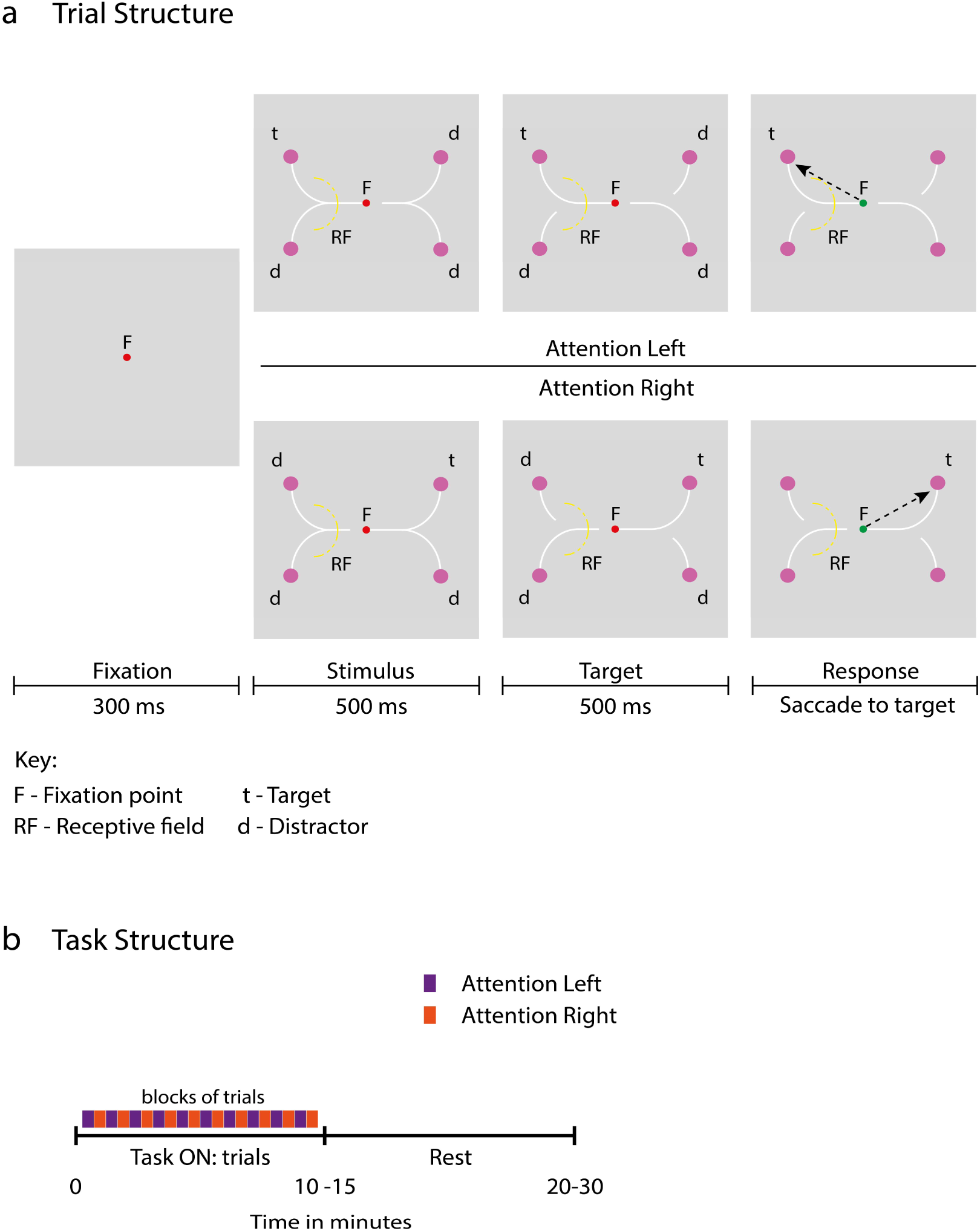
Curve tracing task. (a) The monkey initiates a trial by directing gaze to the red fixation dot (F) for 300ms. Four large purple circles and 4 curves appeared on the screen while the monkey maintained fixation. It was the monkey’s task to make an eye movement to the circle connected by a curve to the fixation point (white target curve) after a delay of 1s. During the first 500ms the fixation point was connected to either the left or right curve but at a second bifurcation both the upper and lower curves were connected so that the entire course of the target curve was not yet revealed. During a second interval of 500ms a contour segment was removed and now the shape of the entire target curve was revealed. Then the fixation dot changed colour to green (Go-cue) and the monkey made a saccade (arrow) to the circle at the end of the target curve, which now was the only circle that was connected to the fixation point circles. The curve tracing task consisted of blocks of 8 trials where the fixation point was either always connected to the curves in the left or the right hemi-field. (b) Periods in which the monkey carried out the task were interleaved with rest periods.

Previous studies demonstrated that humans^34,35^ and monkeys^36^ direct attention to all the segments of the target curve and withdraw attention from the other, distractor curve that is not connected to the fixation point. We aimed to measure acetylcholine release in the macaque dorsolateral prefrontal cortex in monkeys performing a curve tracing task with one target curve and three distractor curves (Fig. 2a,b). Unfortunately, our findings were inconclusive due to the technical challenges that we faced when adapting the electrochemical technique to nonhuman primates. In the discussion, we outline possible improvements that may make the method more viable for future research.

## Methods

### Curve-tracing task

The monkeys performed the task while seated in front of a 21-inch CRT monitor with a refresh rate of 70 Hz and a resolution of 1024 x 768 pixels. The eye position was monitored with a video-based eye tracker (Thomas Recording) and sampled at 250 Hz. The monkey obtained a juice reward at the end of each correct trial.

The behavioural paradigm (Fig. 2a, b) had two conditions, TASK ON and REST. During the TASK ON condition, the monkeys performed a curve tracing task which alternated between blocks ‘attention left’ and ‘attention right’ trials. A trial was initiated when the monkey had maintained his gaze for 300ms within a fixation window, 2.0° in diameter, centered on the fixation point. Four purple circles and four white curves appeared on the screen while the monkey maintained fixation. The fixation point was connected to either the left or right pair of curves so that the monkey knew that he would have to make a saccade into the left or right hemifield. In this phase, the horizontal line elements on either side of the fixation point were still connected to two purple circles so that the monkey did not yet know whether he should make an eye movement to the upper or lower circle. After an interval of 500ms, one contour element at the left and right bifurcation disappeared, hence the entire course of the target curve, connecting the fixation point to one unique purple circle, was revealed. After an additional interval of 500ms, the fixation point changed colour to green, cueing the monkey to make a saccade to the circle at the end of the target curve. The curve tracing task consisted of blocks of 8 trials. During these blocks, the target curve was either always connected to one of the two circles in the left hemi-field (upper panels in Fig. 2a) or to one of the two circles in the right hemi-field (lower panels in Fig. 2a). In other words, the animal either attended the left or right hemifield for blocks of 8 trials. During the REST condition, the monkey did not have to perform any task and the monitor screen was turned off. The Task ON and rest conditions alternated every 10-15 minutes (Fig. 2b).

### Monkeys and surgeries

We carried out the amperometric experiments in an adult macaque monkey (Macaca Mulatta: monkey R), which also participated in other experiments in which we recorded single unit activity (reported elsewhere). We implanted a head post for head stabilization and a recording chamber over the right dorso-lateral prefrontal cortex (dlPFC). During surgeries, general anesthesia was induced with ketamine (15mg/kg injected intramuscularly) and maintained after intubation by ventilation with a mixture of 70% N_2_O and 30% O_2_, supplemented with 0.8% isoflurane, fentanyl (0.005mg/kg intravenously) and midazolam (0.5mg/kg/h intravenously). In a first surgery, the monkey was implanted with a head post. The monkey was then trained on the curve tracing task until he could reliably perform the task. In a second surgery, we performed a craniotomy (centered on stereotaxic coordinates: 21mm anterior, and 17mm lateral) and implanted a titanium chamber (Crist Instruments) for electrophysiological recordings.

All surgical procedures were performed under aseptic conditions and general anesthesia and complied with the US National Institutes of Health Guidelines for the Care and Use of Laboratory Animals and were approved by the Institutional Animal Care and Use Committee of the Royal Netherlands Academy of Arts and Sciences.

### Maintenance of the recording chamber

The chamber was regularly cleaned with Betadine and Hydrogen peroxide. We also used Avastin (Bevacizumab) an angiogenesis inhibitor (slows growth of new blood vessels) in 0.2ml diluted to 1.0 ml saline to decrease the growth of connective tissue over the dura. Avastin helped in keeping the dura clean allowing easier access to the brain tissue^37^. We also removed excess tissue, which grows over the dura mater, at intervals of several weeks, under anesthesia.

We often found it difficult to pass the electrode through the dura and some electrodes broke when we tried to pass them through. We therefore occasionally made a small slit in the dura.

### Preparation of the electrode and solutions

#### Phosphate buffer solution

A 2-litre Phosphate buffer (0.05 M) solution was prepared by dissolving 2.8g of Monosodium Phosphate (NaH_2_PO_4_), 11.34g of Disodium phosphate (Na_2_HPO_4_) and 11.68g of sodium chloride (NaCl) in ddH_2_0. The pH was adjusted to 7.4.

#### m – (1,3) – phenylenediamine

A solution of 5 mM mPD in degassed 0.05 M phosphate buffered saline (PBS). Degassing was accomplished by bubbling nitrogen gas through the 0.05 M PBS solution for 20 minutes to remove oxygen before dissolving mPD. Once the 5 mM mPD (0.045g mPD in 50 mL 0.05 M PBS) dissolved in the degassed 0.05 M PBS, we stored the solution in a brown glass bottle to prevent oxidation under the influence of light.

#### Choline & ascorbic Acid

Stock solution of 20 μM choline (by dissolving 0.07g in 25ml of distilled H_2_0) was used to test the sensitivity of the electrodes during the calibration phase and a stock solution of 20 μM of Ascorbic acid (by dissolving 0.09g in 25ml of distilled H_2_0) was used to test the selectivity of the electrode during the calibration phase. These stock solutions were refreshed every week.

#### Enzyme solution

For the coating of the electrode with enzyme, we prepared a mixture of 1% bovine serum albumin containing 0.01g of BSA and 985 μL distilled H_2_O and added 5 μL glutaraldehyde solution. Choline oxidase solution was diluted as 1 unit/μL and divided into 1 μL aliquots. For the application of the enzyme 1 μL of choline oxidase, 4 μL of BSA-glutaraldehyde solution were mixed together. The presence of bovine serum albumin and glutaraldehyde helps in stabilizing the enzyme.

### Electrode Design

The ceramic-based microelectrode has four 15 x 333 μm^2^ recording sites that are arranged in two pairs (Fig. 1a,b). Every platinum electrode contact was soldered and the electrodes and wires were placed in a scaffold to apply a thick layer of epoxy resin (Max GFE), which cooled and hardened for 24 hours. Then the electrode was removed from the scaffold (Fig. 1c).

We used mPD as an exclusion layer for the electrodes and as substrate for the later immobilization of the enzyme. The layer prevents larger molecules such as ascorbic acid (AA) and dopamine (DA) from reaching the recording surface. Smaller molecules, such peroxide, can pass through the matrix. We placed the electrode in a water bath at 37°C and applied a potential of +0.5 V (Fast Mk II, Quanteon LLC) using an Ag/AgCl indifferent electrode for 20 – 25 minutes to allow the mPD to electropolymerize. The electrode was removed and rinsed with ddH_2_O.

We next coated two of the electrode sites with the enzyme. We applied a mixture containing choline oxidase (CO), bovine serum albumin (BSA) and glutaraldehyde in water (100 nL) to the bottom pair of recording sites using a 1.0-ml syringe (Hamilton, Reno, NV, USA), thus making them sensitive to choline (choline recording channels). The other two recording sites were coated with BSA and glutaraldehyde solution only, and served as sentinel channels. Once the coating had been applied, the microelectrodes were allowed to dry in air at least 24 hours prior to use.

### Recording setup

The recording systems consisted of a potentiostat with 4 channel head-stage (FAST MK II, Quanteon LLC). The recording software was custom designed in Matlab 2009. We measured the signal at a sampling frequency of 20 kHz. To improve signal quality and lower electrical noise, we performed all in-vitro and in-vivo experiments in a Faraday cage. We encapsulated the electrode with a thick layer of two-component epoxy resin for insulation, which improved signal quality and mechanical stability (Fig. 1c).

### Calibration

We calibrated every electrode twice, before and after recording from the dlPFC. We determined the choline sensitivity and the limit of detection (LOD). The electrode was placed in a water bath containing double-distilled water with a magnetic stirrer and we recorded the current, at a holding potential of 0.7 V. The potentiostat keeps this voltage constant so that the reaction of hydrogen peroxide causes a current across the electrode. During the calibration procedure, we first added 5 μl of 20mM of ascorbic acid (AA) to test the quality of the mPD exclusion layer. Electrodes that exhibited a response to AA were not used for recordings in the dlPFC. We then sequentially added 5 μl of 20 mM choline thrice to test the choline-oxidase enzyme complex present on the two recording channels of the electrode. Figure 3a illustrates the traces from an example pre-recording calibration. Ascorbic acid (AA) caused little change in the current, whereas choline increased the current passing through the enzyme channels but not through the non-coated sentinel channels, which remained at baseline (Fig. 3a). The mean choline sensitivity across all electrodes used in the present study during pre-calibration was 16.4 ± 4.7 pA/μM (mean ± s.d.).

**Figure 3.**
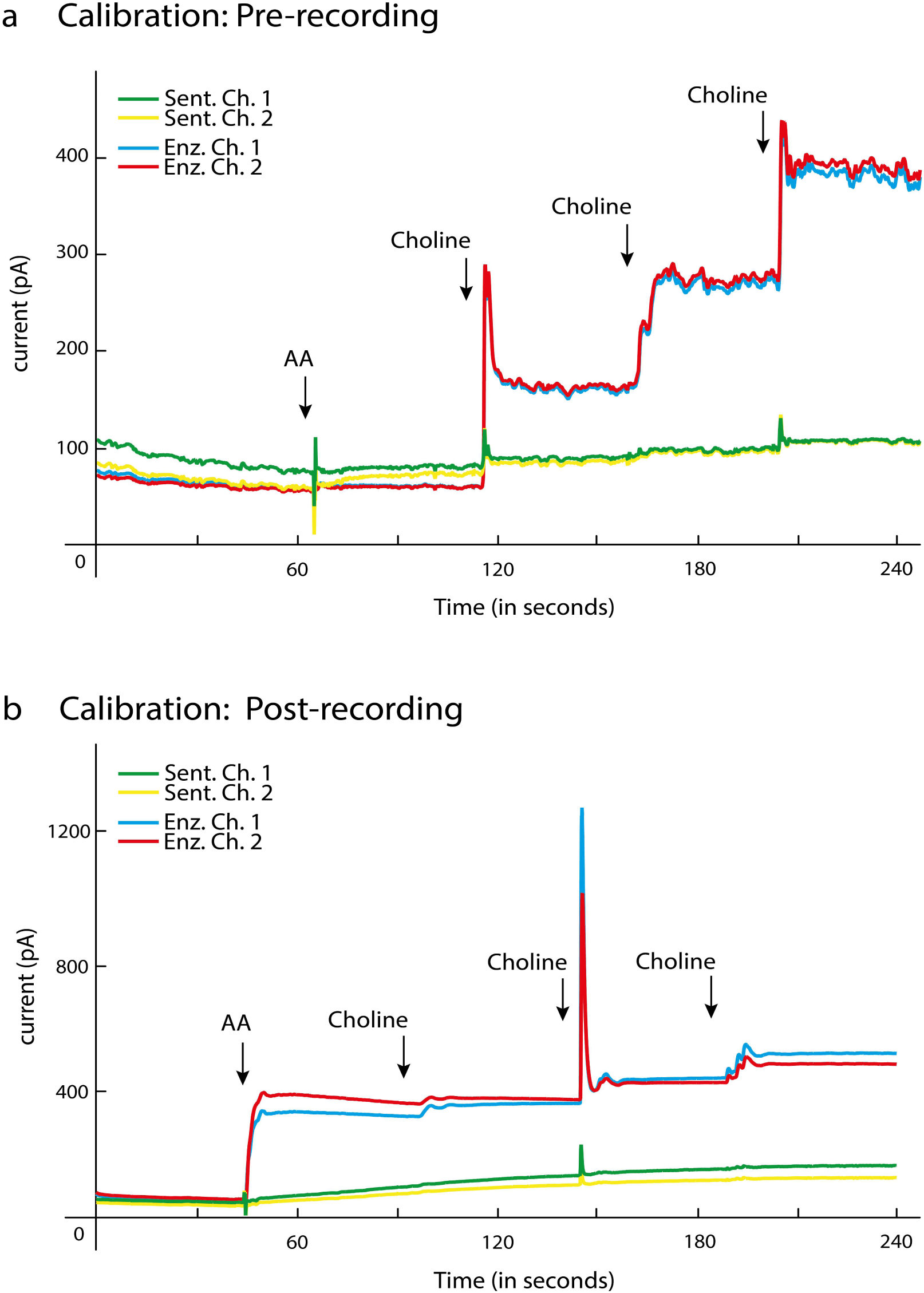
In vitro calibration procedure. In-vitro calibration recordings of the electrode are used to selectivity for choline compared to ascorbic acid (AA). There are four channels, 2 sentinel channels (green and yellow) and 2 enzyme channels (blue and red). (a) Calibration performed before the recording session. The electrode shows a robust response whenever choline is injected into the solution. Note that the electrode is selective for choline because there is no response to AA. (b) Calibration performed after the recording session in the dlPFC. The electrode shows a weakened response to choline and selectivity is lost as it gives a strong response to AA.

We also carried out a post-recording calibration. Figure 3b illustrates the traces of the same electrode as in Figure 3a. We found in this example recording that ascorbic acid (AA) now increased the current for the enzyme-coated channels, which may indicate damage to the mPD exclusion layer. Furthermore, entry into the dlPFC reduced the choline sensitivity of the enzyme-coated electrodes. Across all recordings, the sensitivity decreased from 16.4 ± 4.7 pA/μM pre-recording to 9.2 ± 6.1 pA/μM post-recording (Fig. 4a). This reduction of choline sensitivity was significant (two tailed t-test: t (12) = 5.9, p < 0.001). Figure 4b shows the change in sensitivity for individual recording days.

**Figure 4.**
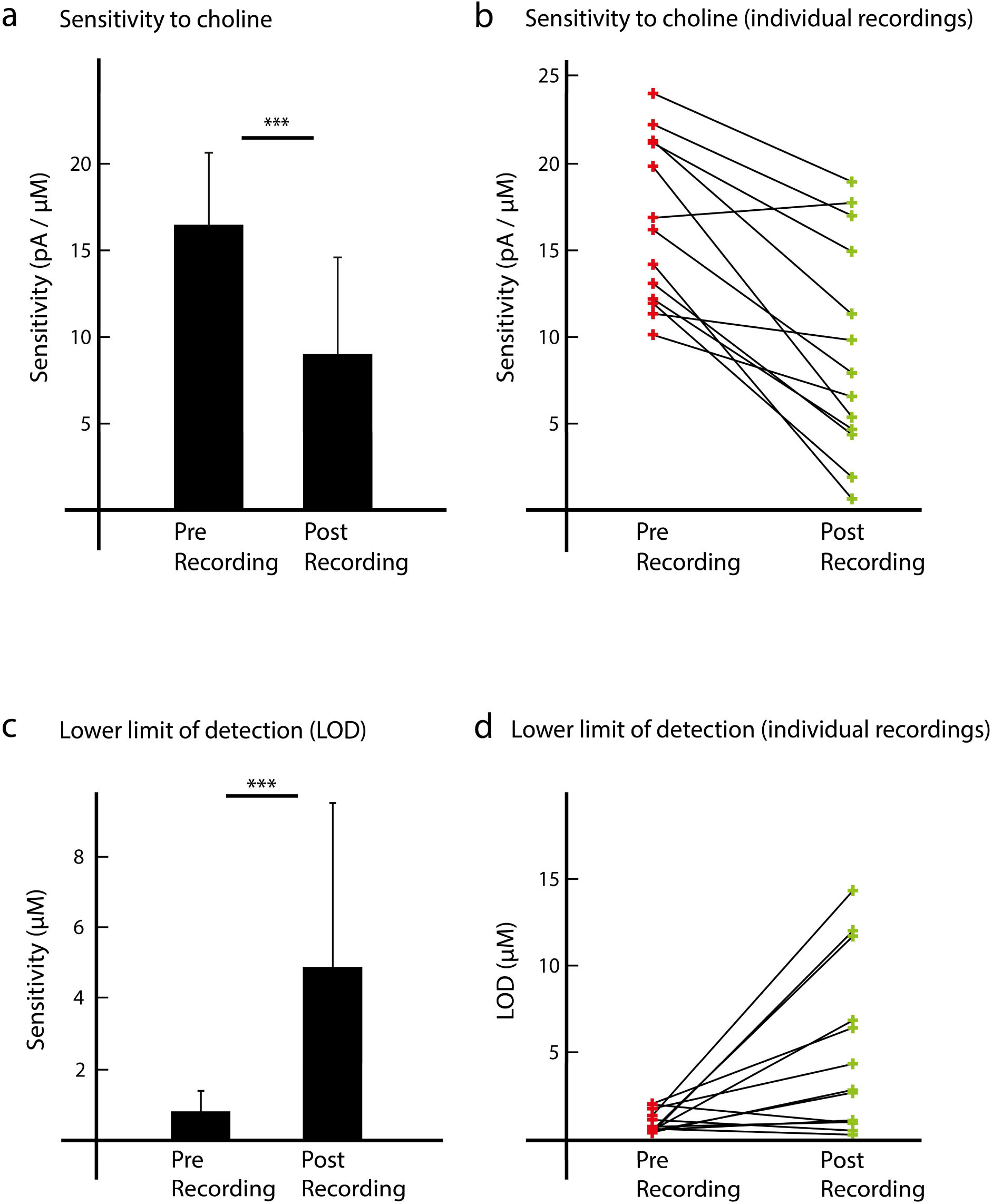
Sensitivity to choline and limit of detection (LOD) of the electrodes. (a) The average sensitivity of the electrode for choline before (pre-recording) and after (post-recording) the experiment. Error bars indicate s.d. (b) The choline sensitivity before and after the experiment for individual recording days. (c) Average limit of detection (LOD) of the electrode for choline before and after the experiment in dlPFC. LOD is a measure of the minimal change in choline concentration that can be detected. (d) LOD before and after the experiment for individual recording days.

We next examined the limit of detection (LOD) of the electrode during the pre- and post-calibration (Fig. 4c). LOD is the lowest quantity of a substance that be detected and we estimated it as three standard deviations of the noise. The mean pre-calibration LOD was 0.80 ± 0.6 μM and the mean post-calibration LOD was 4.8 ± 4.9 μM, a difference that was significant (two-tailed t-test: t (12) = −2.93, p = 0.012). After the recordings, the electrode required a larger amount of choline to register the signal. Figure 4d shows the change in LOD for individual recording days.

## Results

Once the electrode was successfully placed in the cortex, the monkey either engaged in the curve-tracing task or was at rest. The monkeys’ accuracy in the curve-tracing task was always above 75%.

Out of twelve attempts to carry out electrochemical detection of the choline concentration, we only had four recordings in which we recorded changes in the current measured by the potentiostat that appeared to be related to the task. In the other eight recordings, the signals on the enzyme channels and the non-coated sentinel channels were virtually identical. Figure 5a shows the currents across the four channels (two sentinel electrodes and two enzyme-coated electrodes) for one of the more successful recordings. Upon entry of the electrode into the brain, it took about 10 minutes before the currents stabilised as indicated by the rapid and steep descent of the current. This was followed by a more gradual increase in the overall currents that was visible for all four channels. Changes in the concentration of acetylcholine are expected to cause a change in the current across the enzyme-coated electrodes, which is not visible on the non-coated sentinel electrodes.

**Figure 5.**
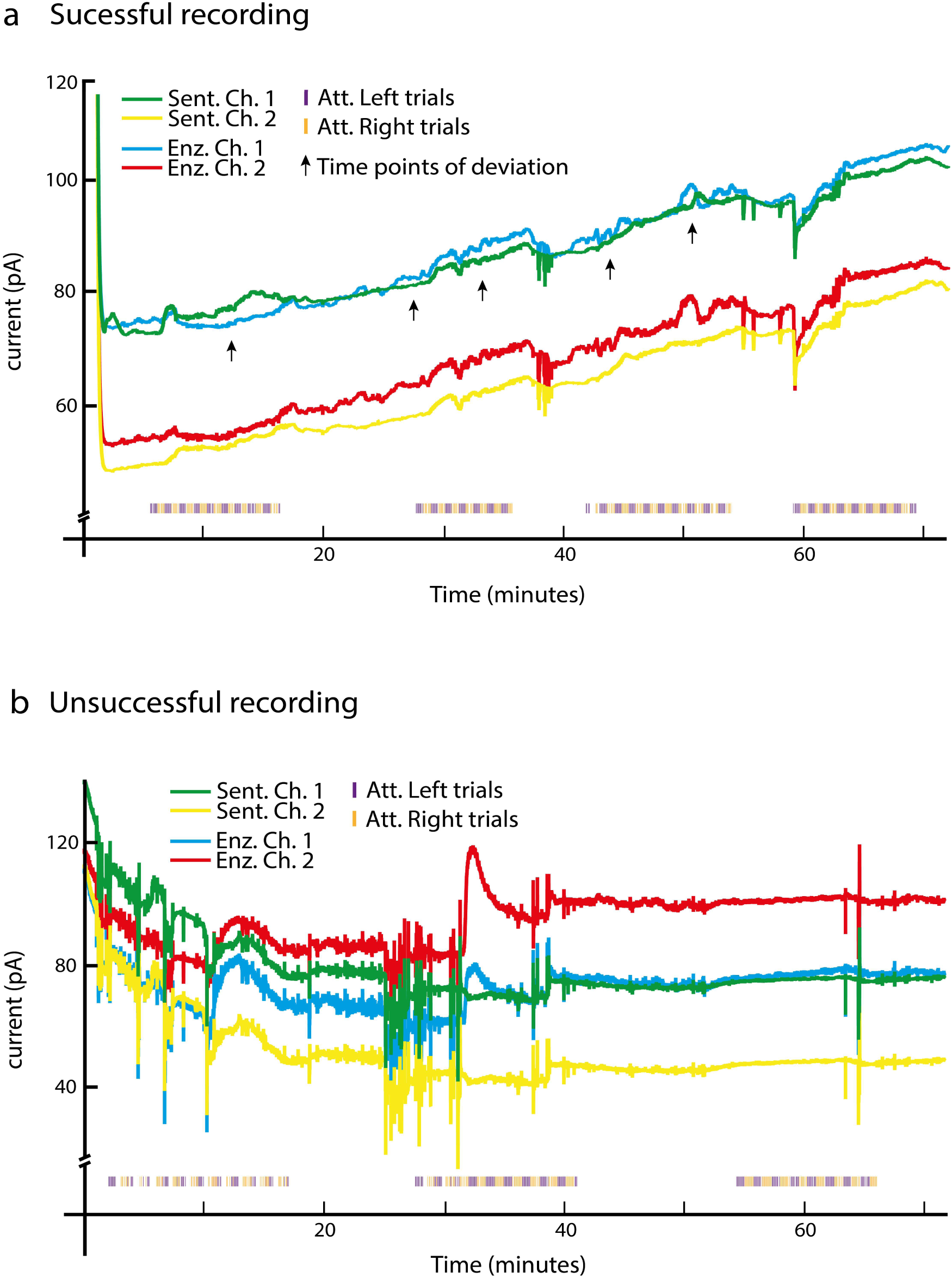
Choline electrochemistry in dlPFC. (a) Successful recording where the signal of the enzyme coated channels (blue and red) showed small deviations from the signal of the sentinel channels (yellow and green). The orange and blue symbols on the x-axis depict the type of trials (attention to the left or right hemi-field). Black arrows illustrate time points where the signal of the enzyme channels shows small fluctuations that differ from those of the sentinel channels. (b) Example unsuccessful recording where activity on the enzyme and sentinel channels was highly similar.

We inspected the data for putative slow changes in the concentration of acetylcholine while the monkey performed the curve tracing task and the REST condition (Fig. 5). These changes in the acetylcholine concentration are expected to cause a difference in the signal between the enzyme-coated channels and the sentinel channels and some of these putative events have been marked with arrows in Fig. 5a. However, we did not observe clear differences between the epoch in which the monkey carried out the curve tracing task and the REST periods in between. In other, unsuccessful penetrations (8 out of 12), the signals from the sentinel and enzyme-coated channels were similar (Fig. 5b). We hypothesized that the absence of differences in the current passing through the enzyme-coated and sentinel channels might have been caused by the enzyme layer shearing off during insertion or by the degradation of the mPD exclusion layer. In both successful and unsuccessful recordings, we observed that the degree of modulation of the current decreased as time progressed. Typically, the fluctuations of the currents across sentinel and enzyme channels became very similar within one hour of recording. We next investigated the signals of the different electrodes at the time-scale of tens of milliseconds. Fig. 6a shows the currents produced by the potentiostat, averaged across blocks of trials in which the monkey directed attention either to the left or right hemifield with activity aligned on stimulus onset (0ms). Interestingly, we saw a clear and reliable modulation of the currents passing through both the enzyme-coated and sentinel electrodes. The circuitry of the potentiostat keeps the potential between the electrodes and the Ag reference electrode constant. We suspect that these relatively fast changing currents of the potentiostat are caused by an influence of evoked potentials in the local EEG, elicited by the onset of the stimulus and the preparation of the eye movement response. The potentiostat tries to maintain the potential at a constant value, and hence it will also compensate for evoked potentials by changing the current. In support of this hypothesis, these current fluctuations occurred at a time-scale of tens of milliseconds, which seems too fast for the diffusion of choline through the different layers of the electrode and the various chemical reactions required by choline-electrochemistry. Accordingly, these fast changes in the currents were equally prominent for the enzyme-coated and sentinel electrodes.

**Figure 6.**
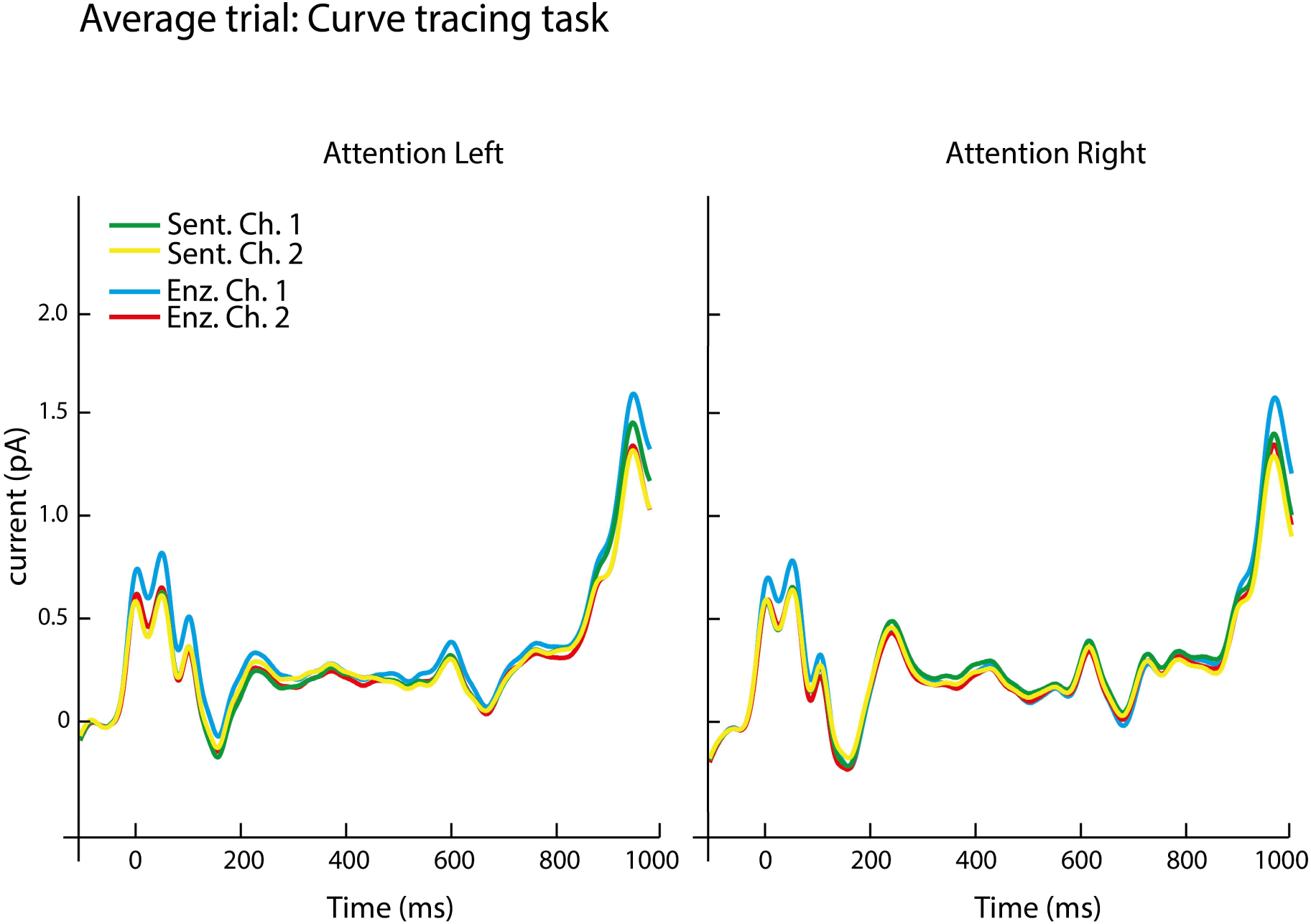
Signal from the potentiostat at a fast time-scale. Average current at the two enzyme-coated (blue and red) and two sentinel channels (yellow and green) for trials with attention directed to the left and right hemi-field. Note the fast signal changes induced by the presentation of the visual stimulus (at time 0) and the preparation of an eye-movement response (at 1000ms).

## Discussion

We aimed to measure the influence of attention on the release of acetylcholine in the dlPFC using a new electrochemical method, but we did not succeed. The influence of attention shifts neuronal activity has been measured in numerous studies and many of them have implicated the cholinergic system in the mechanisms that cause attentional shifts^8,9^. However, the measurement of acetylcholine release at a time-scale of seconds or below has proven to be difficult. While trying to measure these changes in acetylcholine levels in monkey dlPFC, we faced several technical challenges. In spite of our successful in-vitro calibration, only 4 out 12 in-vivo recordings were successful, because there were small differences between the currents at the enzyme-coated and sentinel channels. We did not observe a clear correlation between these possible cholinergic events and the engagement of the monkey in the task. We will now address several issues that will have to be addressed by future studies that aim to apply choline electro-chemistry in the cortex.

Firstly, we observed a degradation of the signal over time. Part of this decrease in the signal was presumably caused by the loss of enzyme activity. We observed a significant reduction in the sensitivity (Fig. 4a,b) and LOD (Fig. 4c,d) during the post recording calibration compared to the pre-recording calibration. This may have been the result of tissue reactions and mechanical damage caused by the penetration of electrode through the dura mater and into the brain^38^.

Secondly, our results also suggest a degradation of the specificity of the signal, because the electrodes started to respond to AA during the post-calibration measurements. This result suggests that the mPD exclusion layer had deteriorated so that the electrodes might also start to respond to variations in the concentration of dopamine, 3,4-Dihydroxyphenylacetic acid (DOPAC) and other substances (Fig. 3b).

### Potential improvements of the method

There is scope for design improvements of the recording chamber which would to make the amperometric recordings more robust. Our results suggest that the mechanical damage to the enzyme coating during electrode penetration across the connective tissue and the dura was an important factor determining the success or failure of the experiment. Jackson & Mithuswamy^39^ have detailed the use of a transparent soft silicone gel covering the cortex of rodents as a method to safely insert electrodes multiple times. However, this method requires removal of the dura matter and insertion of the silicone gel and it is unknown if this method is safe for the monkey cortex. Other studies have explored the use of artificial dura for recording chambers with an improved design for attachment^40^.

Another improvement which can be made is in the design of the electrode. Figure 7 shows a schematic of proposed design changes to the electrode. In the new design the platinum contact points are places inside ‘wells’. These wells can be filled with the enzyme complex and could protect the coatings from being sheared off during the insertion into the cortex. Another improvement could be made at the tip of the electrode. If the tip is made out of metal or a hard plastic material and its shape/profile is sharper, it will penetrate more easily through the dura.

**Figure 7.**
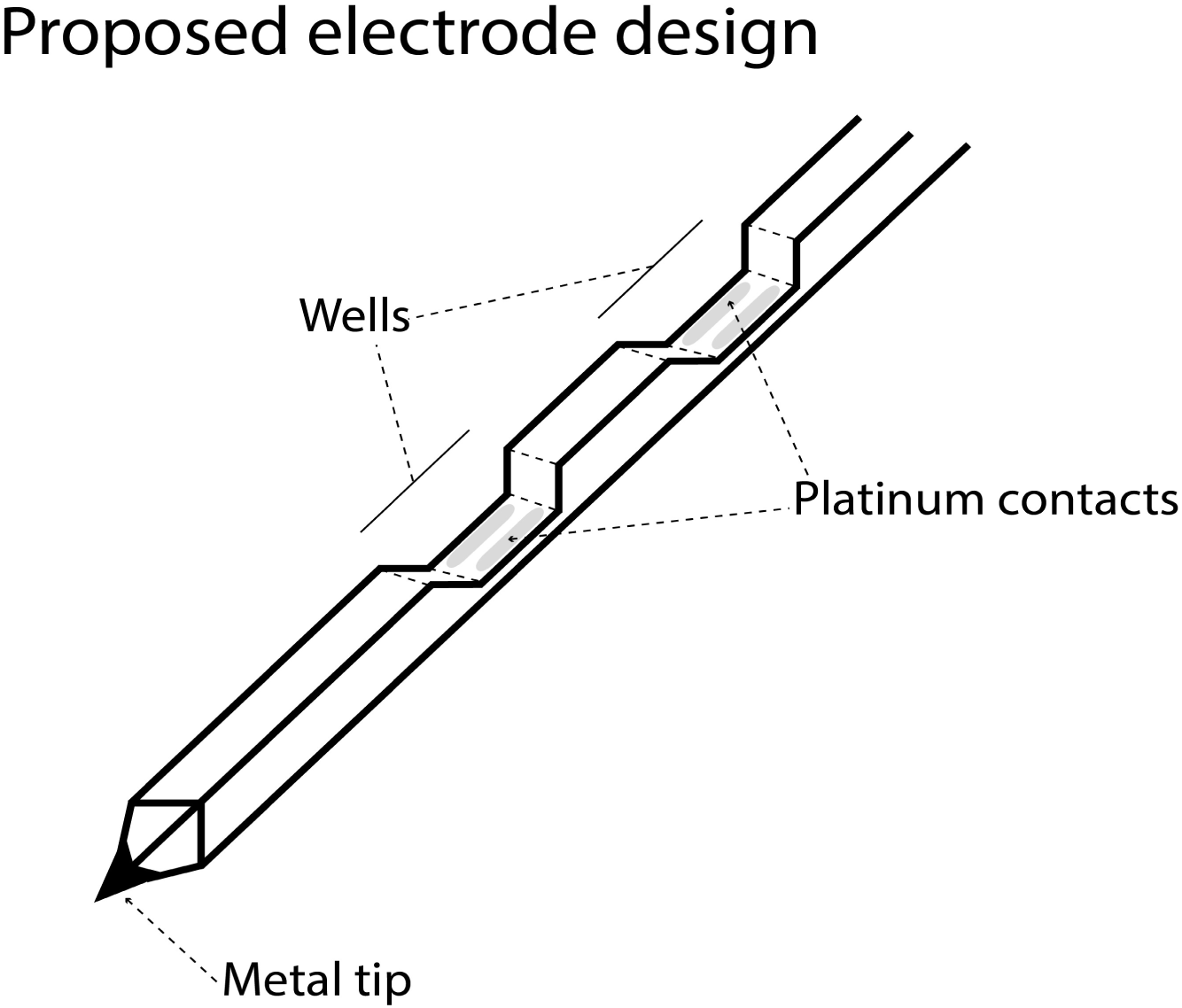
Proposed future electrode design with improved mechanical qualities. Deposition of the mPD and enzyme layers in a cavity of the electrode might improve the mechanical stability of these layers. A sharper tip would facilitate penetration of the dura mater.

In the recent years, there have been many advances in polymer material technology^41^. For example, conductive polymers such as Polyaniline (PANI), poly3,4-ethylenedioxythiophene (PEDOT) have been created, which are biocompatible and might in the future be adapted for use as biosensors. These polymers are also chemically and thermally stable and can bind enzymes. Other advances in nanotechnology have resulted in sensors for acetylcholine using carbon nanotubes and acetylcholinesterase thin films^42^, but their application has so far been restricted to in-vitro setups. The transition from in-vitro to in-vivo systems can be challenging and other technical issues may arise when using these methods in the brain.

In this study, we have outlined the problems that we faced when adapting choline electrochemistry to non-human primates. We have outlined a few potential solutions and improvements that will hopefully allow future scientists to develop better and suitable sensors for the release of neuromodulatory substances, such as acetylcholine, during cognitive tasks.

## Acknowledgments

This work was supported by EU (Marie-Curie Action PITN-GA-2011-290011 awarded to PRR.

## Author contributions

DV, CvdT, BvV and PRR conceived and designed the experiments. DV and BvV collected the data. DV analyzed the data with advice from CvdT and PRR. DV and PRR wrote the paper. All authors reviewed the manuscript.

